# Killer Toxin K28 resistance in yeast relies on COG complex mediated trafficking of the defence factor Ktd1

**DOI:** 10.1101/2024.12.20.629825

**Authors:** Kamilla ME. Laidlaw, Hatwan H. Nadir, Amy Milburn, Martha S.C. Xelhuantzi, Justas Stanislovas, Alastair P. Droop, Sandy MacDonald, Ilya Andreev, Andrew Leech, Daniel Ungar, Meru J. Sadhu, Chris MacDonald

**Affiliations:** York Biomedical Research Institute and Department of Biology, University of York, York, UK; Bioscience Technology Facility, Department of Biology, University of York, UK; Centre for Genomics & Data Science Research, National Human Genome Research Institute, NIH, Bethesda, MD

## Abstract

A/B toxins are a diverse family of protein toxins that enter host cells via endocytosis and induce cell death. In yeast, the A/B toxin K28 is internalised to endosomes of susceptible yeast, before following the retrograde trafficking pathway and ultimately triggering cell cycle arrest. The endolysosomal defence factor Ktd1 protects against K28, but its regulation remains unclear. Cog7, a subunit of the conserved oligomeric Golgi (COG) tethering complex, has been implicated in K28 defence, though the mechanism is unknown. We developed a high throughput K28 sensitivity assay and bespoke analysis package to show that all lobe B COG subunits (Cog5 - 8) are required for K28 resistance. Although the COG complex modulates glycosylation of the surface molecules required to bind extracellular K28, our experiments reveal that the hypersensitivity of *cog* mutants is primarily explained by defects in Ktd1 trafficking. Ktd1 mis-localisation in *cog* mutants is reminiscent to disruptions in Snc1, a surface cargo that recycles multiple times via the Golgi. This work suggests not only that the COG complex is responsible for the precise trafficking Ktd1 required to mediate toxin defence, but that Ktd1 may survey endolysosomal compartments for internalised K28. This work underpins the importance of Ktd1 in defence against the A/B toxin K28, and implies various membrane trafficking regulators might influence toxin effects in other eukaryotic systems.

## INTRODUCTION

Protein toxins are molecular weapons secreted by cells to target other organisms, playing a key role in evolutionary processes (Stern, 2013). Toxins of the A/B family, which includes cholera, Shiga, diphtheria, and ricin, are secreted by diverse organisms and target a variety of host cells to initiate cell death, through different modes of action (Odumosu et al., 2010). Despite this, most A/B toxins also share various features, such as a bipartite structure consisting of two moieties: an enzymatic active A subunit and a B subunit responsible for binding host cells, although subunit stoichiometries can vary (Márquez-López and Fanarraga, 2023). A/B toxins bind host cells and enter via endocytosis, where the toxin molecules are internalised to endosome compartments before trafficking through the retrograde pathway and triggering cell death (Falnes and Sandvig, 2000).

The yeast Killer Toxin 28 (K28) is a member of the A/B toxin family that is produced by yeast cells infected with the dsRNA M28 virus (Schmitt and Tipper, 1990). K28 is synthesised as a pre-protoxin, and is modified and processed during trafficking through the secretory pathway *en route* to the plasma membrane for secretion (Breinig et al., 2006). Secreted K28 toxin binds to glycosylated residues on the surface of host cells and then enters the cell using canonical endocytic machinery (Eisfeld et al., 2000; Schmitt and Radler, 1987). Internalised K28 traffics from endosomes through the retrograde pathway, ultimately translocating to the nucleus via the cytoplasm to induce G1/S cell cycle arrest (Schmitt et al., 1996). Ktd1 (Killer toxin defence factor 1) is a membrane protein that localises to the endolysosomal system in yeast and provides resistance to K28 (Andreev et al., 2023). Ktd1 is a member of the Dup240 gene family, members of which have two transmembrane domains (Poirey et al., 2002). Ktd1 mainly localises to the limiting membrane of the vacuole, but is also found in other endolysosomal compartments (Andreev et al., 2023). Despite its critical role in K28 defence, the mechanism by which Ktd1 protects cells from toxin remains unknown.

The conserved oligomeric Golgi (COG) complex is a conserved multi-subunit complex that is involved in tethering retrograde vesicles at the Golgi through coordination of soluble NSF (N-ethylmaleimide sensitive factor) attachment protein (SNAP) receptor (SNARE) mediated membrane fusion (Willett et al., 2013a). As such, the COG complex is responsible for organisation of Golgi localised glycosylation enzymes that modify transiting glycoproteins. Defects in this processing are associated with a variety of human diseases, including congenital disorders of glycosylation and neurological disorders (Foulquier, 2009). The yeast COG complex is composed of eight subunits (Cog1 - 8) that form two subcomplexes, Cog1 - 4 (lobe A) and Cog5 - 8 (lobe B) and is related to other multi-subunit complexes (Fotso et al., 2005; Szentgyörgyi and Spang, 2023; Whyte and Munro, 2002). One subunit of the COG complex has been implicated in K28 sensitivity, as a genetic screen for mutants with altered response to K28 revealed that *cog7Δ* cells are hypersensitive (Carroll et al., 2009). Whether the role of Cog7 is related to maintenance of the Golgi structure, proper modification of glycosylated K28 binding surface proteins, coordination of proteins transiting the Golgi, or an alternative mechanism is not known. Intriguingly, the genetic screen that implicated Cog7 in K28 defence tested various other mutants of the COG complex and did not find any with altered sensitivity to toxin.

In this study we explore the role of the COG complex in K28 resistance, describe a more sensitive and higher throughput assay to measure growth of yeast exposed to media ± K28 toxin, and propose a model for Ktd1-medated defence that relies on precise membrane trafficking to endolysosomes for toxin capture and neutralisation.

## RESULTS

### The COG complex is implicated in K28 toxin defence

A previous genetic screen for killer toxin K28 sensitive and resistant mutants (**Figure 1A**) was performed using a 96-well based ‘halo’ assay from a spot of killer toxin secreting cells placed on a lawn of tester yeast (Carroll et al., 2009). This screen identified *cog7Δ* mutants as hypersensitive to K28. To test whether this effect was specific to mutations in *COG7* or more generally through abrogation of the COG complex, we validated deletion mutants of the non-essential lobe B subunits: *cog5Δ, cog6Δ, cog7Δ, cog8Δ* (**Figure 1B**). Mutants of lobe B have previously been reported to exhibit a growth defect at elevated temperature (Gao and Banfield, 2020; Sinha et al., 2008). Lobe B *cog* mutants were shown to grow poorly at 37°C when compared to wild-type (WT) cells (**Figure 1C**), although this effect was only observed when cells were grown on rich media, with no obvious differences in synthetic complete (SC) minimal media (**Figure 1D**). Although wild-type cells have reduced ability to grow when at elevated temperature on media with alkaline pH, we find this combination is particularly detrimental for growth of the non-essential COG complex mutants. As a control for perturbation of glycosylation, no defects were observed at elevated temperature for *mnn2Δ* mutants, which lack the Golgi resident alpha-1,2-mannosyltransferase Mnn2 (Raschke et al., 1973; Rayner and Munro, 1998).

**Figure 1:**
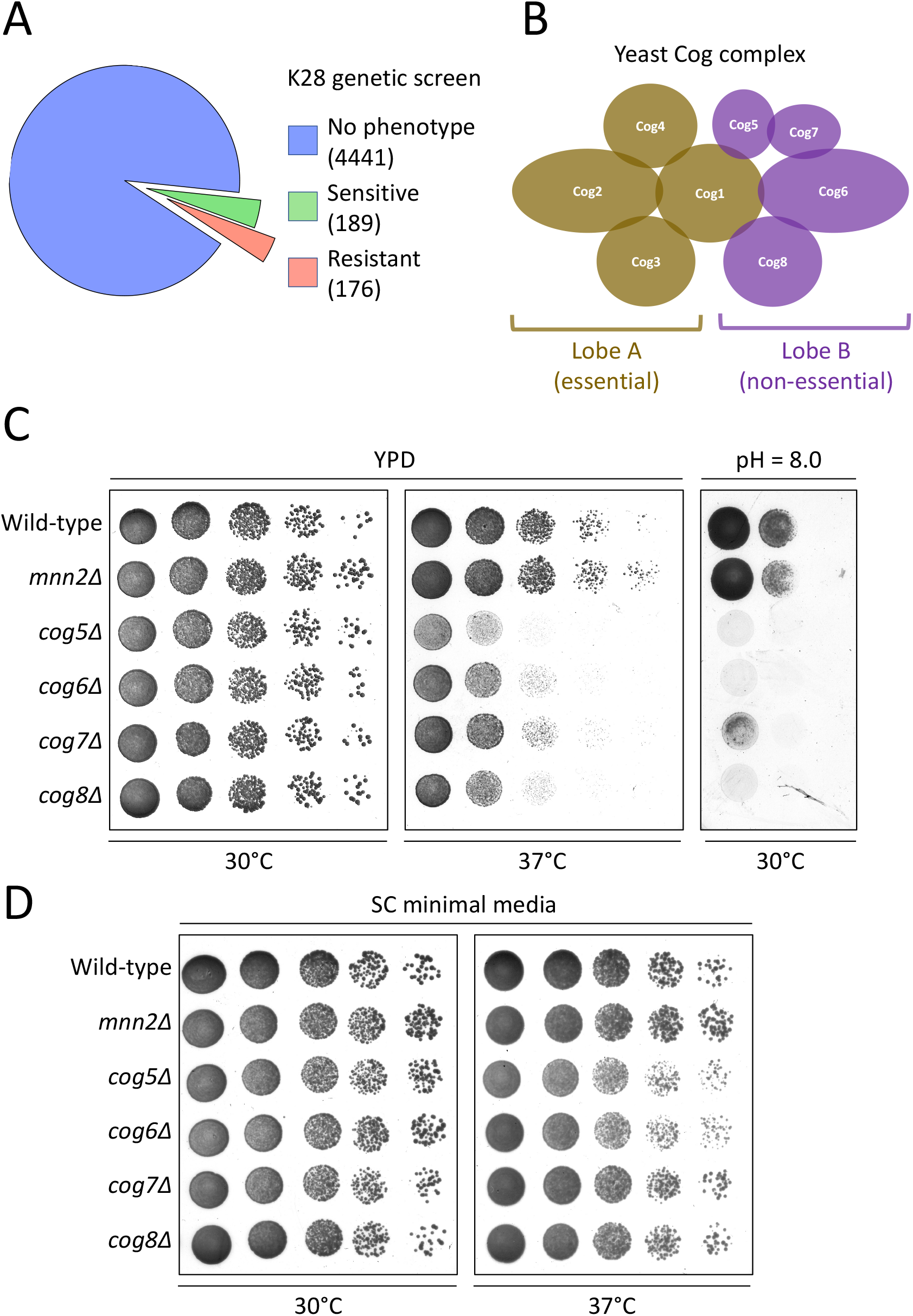
Deletion of non-essential COG complex subunits causes growth defect. **A)** Summary of K28 sensitivity genetic screen performed by (Carroll et al., 2009), showing mutants tested that have no change upon K28 treatment (blue), or those that are either hypersensitive (green) or resistant (red). **B)** Schematic representation of the yeast Cog complex comprised of 8 subunits organised between Lobe A (Cog1 - 4; yellow) and Lobe B (Cog5 - 8; purple), with some of the main physical interactions depicted with subunit overlap (Fotso et al., 2005). **C - D)** Wild-type (WT) and indicated mutant strains were grown to mid-log phase overnight in YPD media before normalised cell pellets were harvested and diluted 10-fold in water prior to 5μl spotted onto solid agar media: YPD **(C)**, or SC **(D)** and incubated at either 30°C or 37°C prior to recording growth.

The halo phenotype generated on a lawn of yeast cultured in the presence of K28 secreting cells has a relatively low dynamic range. Before assessing K28 sensitivity of non-essential *cog* mutants we first optimised the halo assay for maximum halo size produced from *ktd1Δ* cells, which are hypersensitive due to the loss of the Ktd1 defence factor (Andreev et al., 2023). High density of cultures used to create lawns correlates with reduced halo size, with the optimal OD_600_ = 0.1 - 0.2 (**Figure 2A**). Next, the culturing media, of both the lawn but also the K28 secretor strain, were tested for effects on halo radius. For this, WT (BY4741 and BY4742) and hypersensitive (*ktd1Δ*) lawn strains were grown in rich (YPD) and minimal (SC) media, with and without gelatin, a stabilisation component included in some media recipes for K28 sensitivity (Andreev et al., 2023). Although the difference between media with and without gelatin was not significant, we consistently observed lawns grown in minimal media had larger halos than rich media (**Figure 2B**). Oppositely, we find the K28 secretor strain produces larger halos when grown in YPD. We used these optimised conditions to test K28 sensitivity of lobe B *cog* mutants. As a control for toxin sensitivity, alongside WT cells, we also include *mnn2Δ*, which are resistant to K28 due to reduced binding of K28 (Andreev et al., 2023; Carroll et al., 2009). This assay showed that *cog5Δ, cog6Δ, cog7Δ*, and *cog8Δ* mutants exhibit a significant increase in K28 sensitivity (**Figure 2C**), with all mutants showing similar levels of sensitivity. This suggests that beyond the additional finding that Cog7 is involved in K28 sensitivity, that a general functionality of the COG complex is required for maintaining resistance to K28 toxicity.

**Figure 2:**
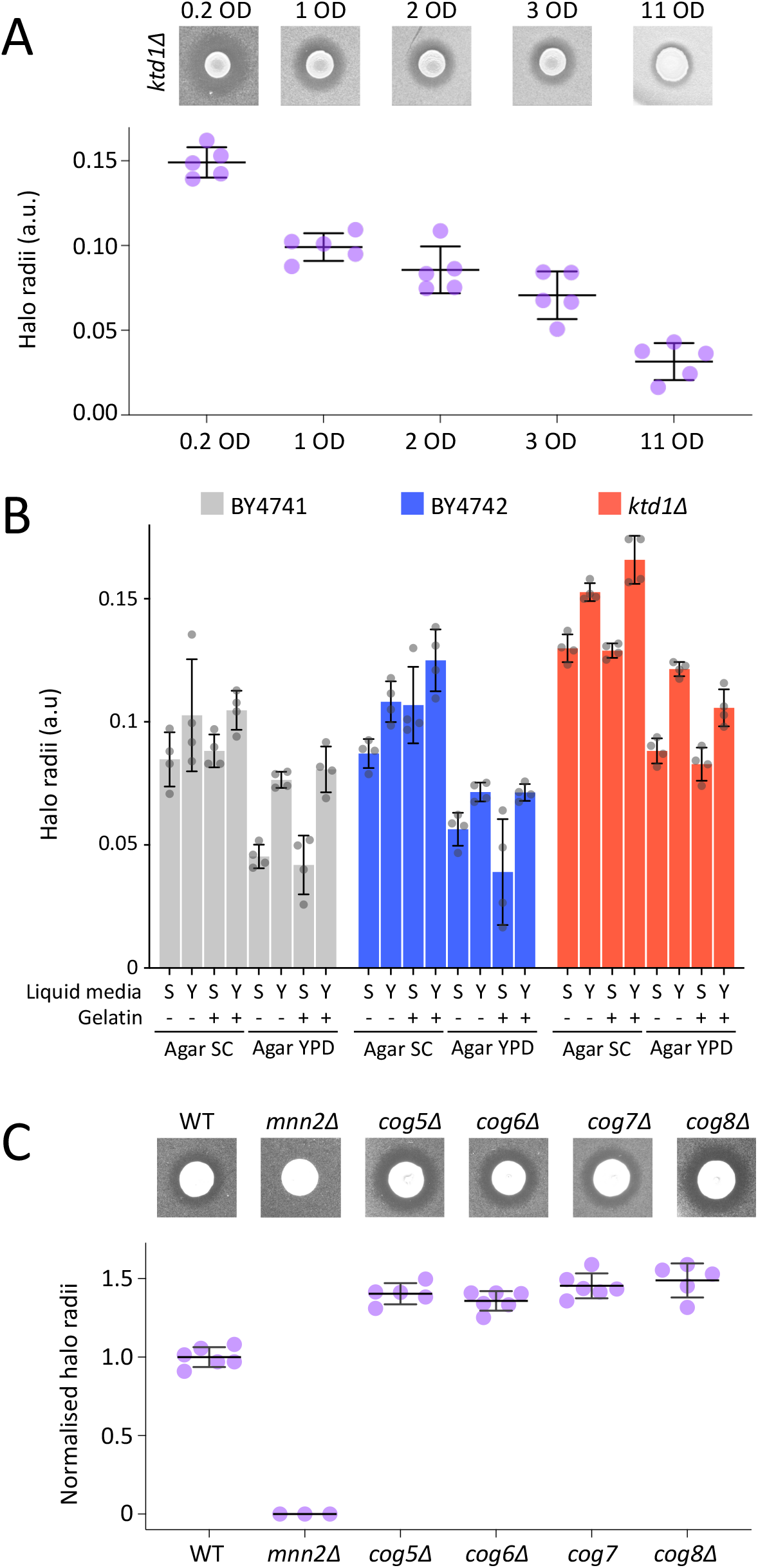
Optimised halo assay shows the COG complex is required for K28 toxin resistance. **A)** Indicated yeast strains were grown to mid-log phase in SC media before lawns at indicated optical densities were created and spread on SC pH 4.7 plates before K28 secretor cells added, with representative halos (upper) and quantified biological repeats (n=5) plotted (lower). **B)** Data from a large series of K28 sensitivity halo assays for two wild-type strains (BY4741 and BY4742) in addition to *ktd1Δ* mutants are plotted. Strains for testing were plated on agar containing either SC or YPD media, with and without gelatin for toxin stability. Comparison of liquid SC (S) or YPD (Y) media used to culture the K28 hypersecretor cells before they were added to lawns was also included. **C - D)** K28 sensitivity halo assays were performed with indicated strains with quantified halo radii (n > 3) plotted below. Error bars show standard deviation.

As COG subunits from lobe A are required for viability (Blackburn et al., 2019; Giaever et al., 2002; Whyte and Munro, 2001) deletion of these genes could not be tested. In an effort to include these COG subunits in downstream analyses, hypomorphic alleles of lobe A subunits were tested. One method to achieve decreased expression of essential genes is the insertion of a cassette in the 3’ untranslated region (UTR), termed Decreased Abundance by mRNA Perturbation (DAmP), which destabilizes the mRNA (Breslow et al., 2008; Schuldiner et al., 2005). However, strains expressing DAmP alleles of both *cog2* and *cog4* were tested but fail to phenocopy lobe B *cog* null growth defects at 37°C (**Supplemental Figure S1A**). Another approach to reduce expression of essential yeast genes is to exchange the endogenous promoter with a tightly regulated estradiol inducible promoter, termed Yeast Estradiol strains with Titratable Induction (YETI) (Arita et al., 2021). However, growing YETI controlled *COG2, COG3* and *COG4* strains in the absence of the estradiol inducer, to minimize expression levels, supported efficient growth at 37°C. However, the lobe A *cog* alleles with reduced levels did not result in K28 sensitivity (**Supplemental Figure S1B**). We conclude that the reduced expression of essential lobe A *cog* mutants does not reduce cellular levels of the COG complex sufficiently to inhibit its capacity to tolerate heat stress or K28 exposure. Therefore, we assume these mutants are not suitable for testing toxin sensitivity of COG lobe A mutant yeast.

### An improved method confirms K28 hypersensitivity to COG lobe B mutants

The low sensitivity of the K28 halo assay might explain why only one of four tested *cog* mutants represented in the deletion libraries were identified in a systematic screen (Carroll et al., 2009). Even with optimisation there is relatively little difference between halo radii from wild-type, resistant and sensitive strains. We have recently optimised a liquid plate growth assay that allows more subtle phenotypes to be observed (Andreev et al., 2023). This assay cultures cells in a microtiter plate and monitors growth over time in media enriched with K28 from a hypersecretor strain, compared with control media from a heat cured strain. To analyse data generated from these K28 sensitivity-based growth assays, we have created a bioinformatic analysis pipeline for 96 well plate liquid cultures to document growth curves (**Figure 3A**). This optimised R:Shiny app is available at https://shiny.york.ac.uk/bioltf/cm_growthcurves/. Data comparisons between strains and across different conditions, such as K28 presence are generated, in addition to statistical analyses (**Figure 3B - 3C, Supplemental Figure S2**). This provides a sensitive and high-throughput analysis pipeline for these K28 assays and could be used for a wide array of applications. We do note that highly resistant strains like mutants with abrogated glycosylation profiles (*mnn1Δ, mnn2Δ, mnn9Δ*) do exhibit very obvious phenotypes with the halo assay due to essentially having zero growth inhibition (**Figure 3D - 3E**). The high throughput liquid plate assay shows the mannosyltransferase null strains grow very similar to wild-type cells, in media with and without K28 (**Figure 3F - 3H**). We assume that wild-type cells expressing *KTD1* sufficiently tolerate K28 to reach near maximal doubling time at exponential phase, showing little difference to the truly resistant (*mnn1Δ, mnn2Δ, mnn9Δ*) strains.

**Figure 3:**
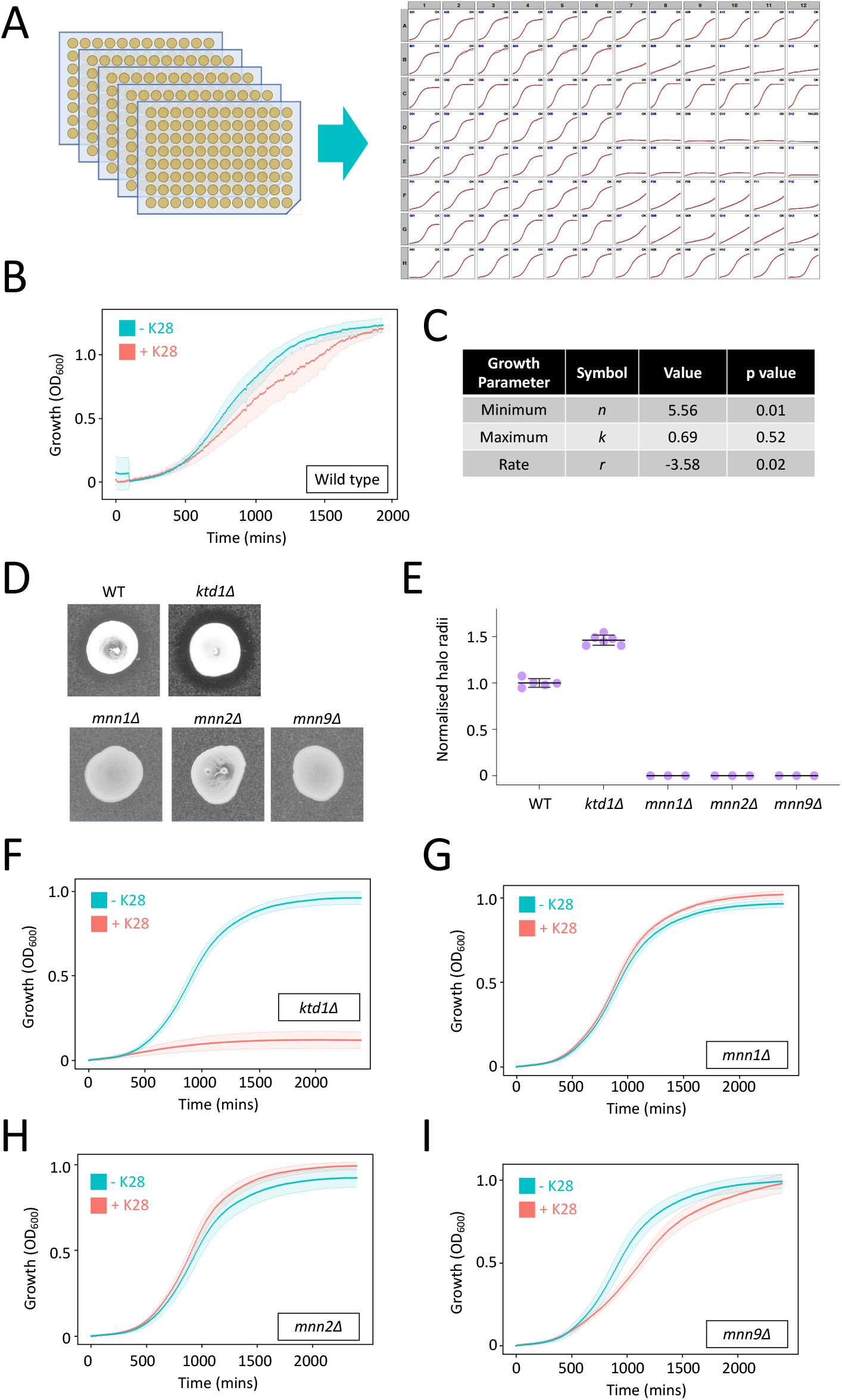
Analysis pipeline for high-throughput liquid assay. **A)** A high throughput 96 well plate liquid assay for yeast growth including example plot used for quality control assessment of growth profiles. **B)** Growth profile of wild-type cells exposed to media -K28 (blue) or +K28, (red) with standard deviation shown in shaded regions. **C)** Growth parameters extracted from the R:shiny app used to generate graphs from raw data imported. **D)** Halo assays of indicated lawns exposed to spot of K28 secretor cells, with quantification (n ≥ 3) depicted. **E)** Quantification from (D). Error bars show standard deviation. **F-I)** Liquid growth assays of indicated cells grown in control media (blue) or K28 enriched media (red).

We conclude that both assays have merits, with the halo and liquid based assays being better for assessing resistance and sensitivity phenotypes, respectively. We then used the liquid assay to measure sensitivity of *cog5Δ, cog6Δ, cog7Δ, cog8Δ* mutants. In agreement with the halo-based previous experiments, all lobe B *cog* mutants grow at very similar rates to wild-type cells over 32 hours in control media lacking K28, but show a significant sensitivity in media containing K28 (**Figure 4A - 4D**). This further supports the notion that a function related to the COG complex, and not an individual subunit, is required for K28 defence.

**Figure 4:**
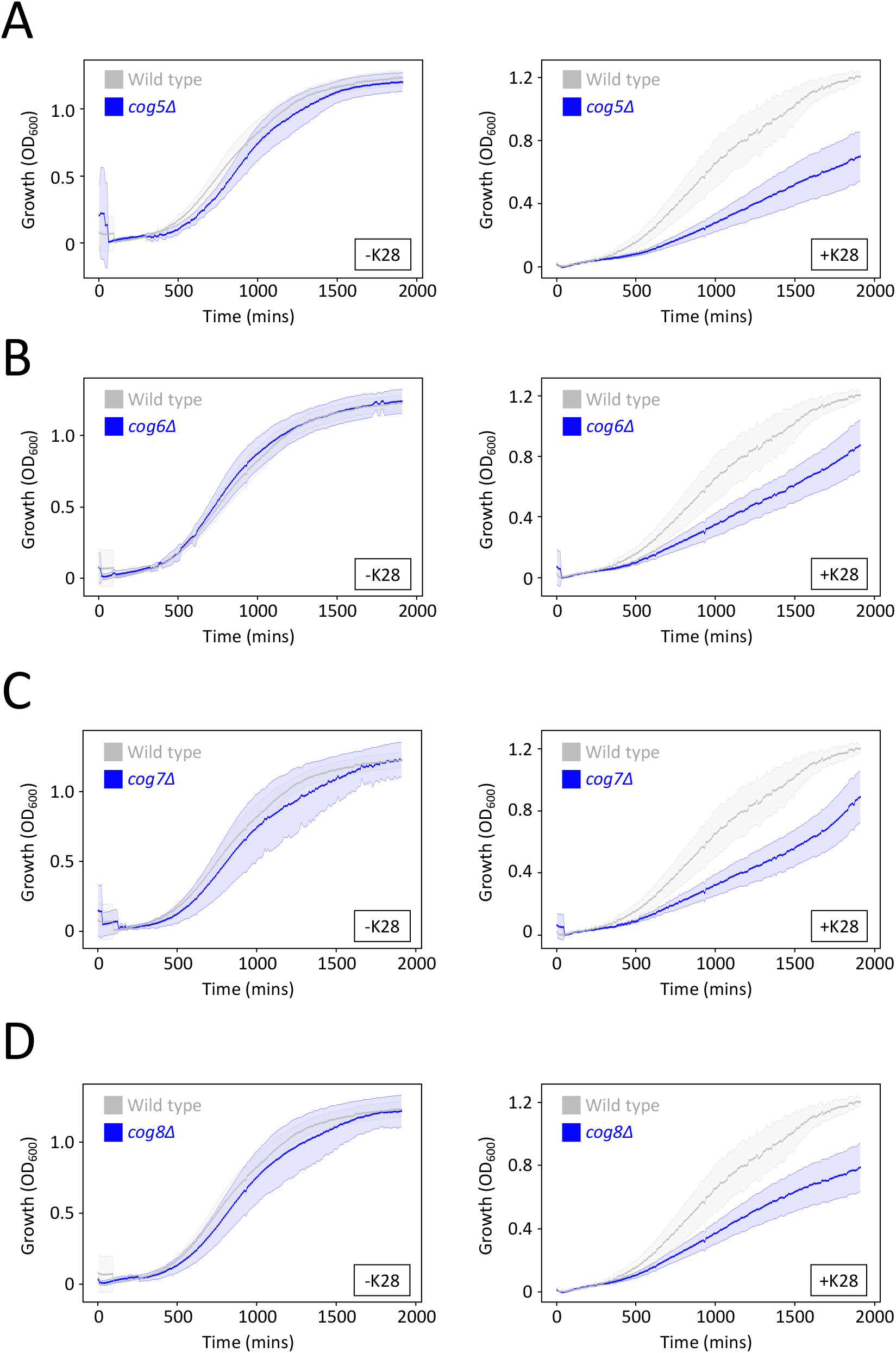
*cog* mutants are hypersensitive to K28 in liquid growth assay. **A - D)** Indicated *cog* mutants (blue) were compared to wild-type cells (grew) grown in same conditions in the same plate, both in media from heat cured yeast (left) and media enriched with active toxin (right).

### K28 binding is unaffected in *cog* mutants

These results suggest that the COG complex is required for defence against K28, but the mechanism is unclear. As in all other eukaryotes, the COG complex in yeast regulates glycosylation of proteins that traffic through the secretory pathway (Fotso et al., 2005; Suvorova et al., 2002; Whyte and Munro, 2001). As mentioned, yeast cells lacking mannosyltransferases, like *mnn2Δ* mutants, are resistant to toxin through reduced capacity for the toxin to bind and enter the cell (Carroll et al., 2009). We hypothesised that deregulation of Golgi mannosyltransferases in *cog* mutants might alter their surface glycosylation profile to increase toxin binding, and therefore explain K28 hypersensitivity (**Figure 5A**). To test this hypothesis, we first created a series of double mutants combining non-essential *cogΔ* deletions with *mnn9Δ*, which lacks the early acting alpha-1,6-mannosyltransferase involved in protein *N*-linked glycosylation (Gopal and Ballou, 1987; Yip et al., 1994). These mutants are resistant to K28, excluding the possibility that an indirect function of the COG complex explains the K28 sensitive phenotype (**Figure 5B**). We next performed adsorption assays to analyse how well K28 bound to WT and *cog* mutants (**Figure 5C**). This data included a *mnn2Δ* control that does not efficiently bind K28. These experiments showed no significant changes in the ability of K28 to bind *cog* mutant cells (**Figure 5D**). We predicted that adsorption of K28 in all four lobe B *cog* mutants would need to be obvious to explain their high sensitivity to K28 phenotype.

**Figure 5:**
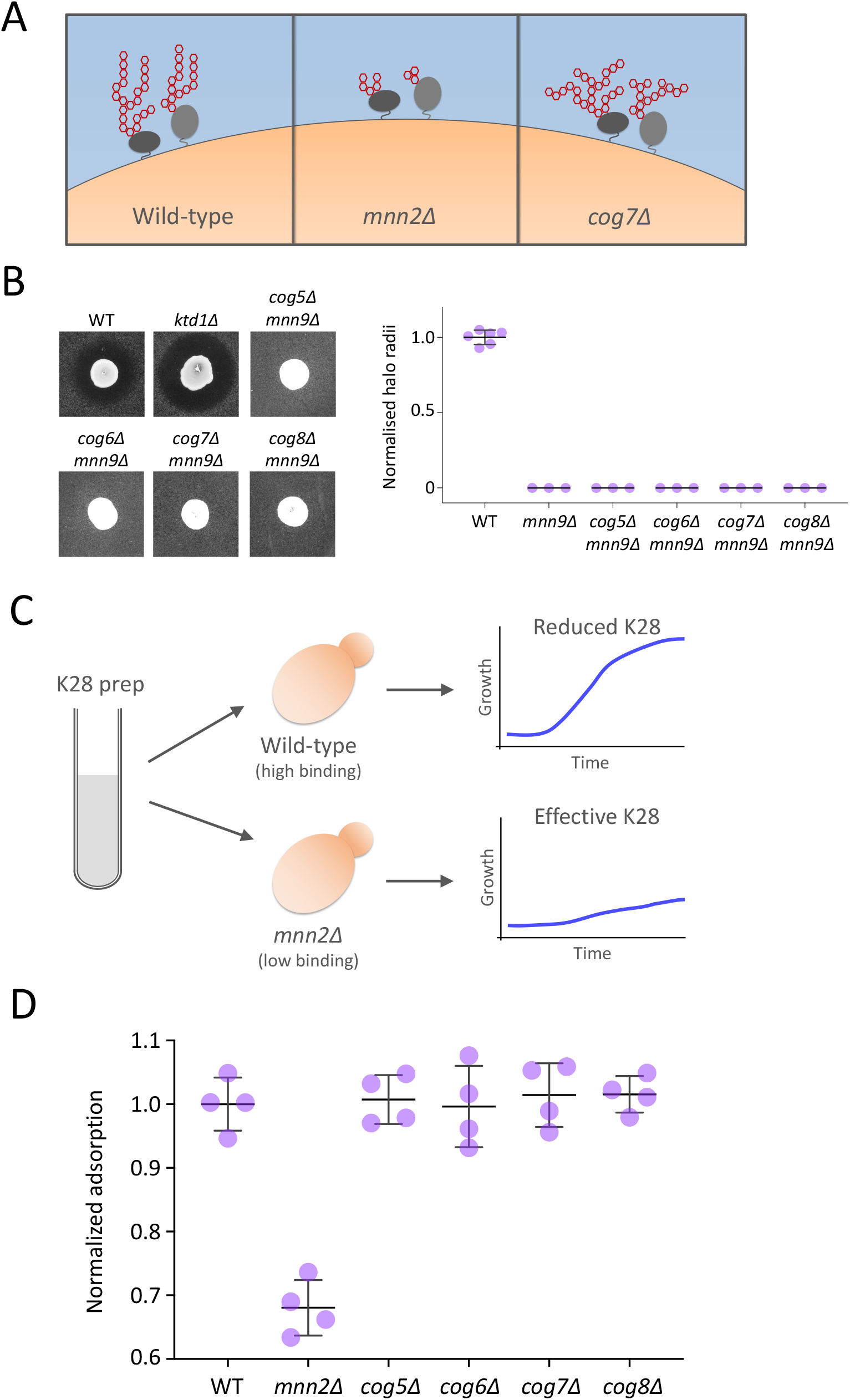
Hypothetical model for COG complex-mediated K28 binding. **A)** Cartoon model to depict hypothesis that mannosyltransferase mutants like *mnn2Δ* have reduced binding of K28 but that *cog7Δ* cells could have altered glycosylation profiles that have higher toxin affinity. **B)** K28 based halo assays for indicated mutants, including quantifications (n ≥ 3). **C)** Basis for adsorption assay, where media enriched for toxin is preincubated with tester cells for adsorption, before cells removed and toxin efficacy in media is tested indirectly by growth capacity of wild-type cells. **D)** Adsorption assay results normalised to wild-type cells and compared across indicated strains (n = 4). Error bars show standard deviation.

### The COG complex is required for proper vacuolar sorting of Ktd1

As elevated adsorption does not explain *cog* mutant K28 sensitivity, we explored other explanations. Although the COG complex does coordinate Golgi enzymes, it achieves this through a fundamental regulation of membrane trafficking at the Golgi (Miller et al., 2013; Willett et al., 2013a; Willett et al., 2013b; Zolov and Lupashin, 2005). We therefore set out to test the idea that Ktd1 is mislocalised in *cog* mutants. To facilitate expression of GFP tagged Ktd1 in many strains, we created a plasmid that expresses GFP-Ktd1 from the *CUP1* promoter. In media lacking copper, we find this construct expresses to similar levels and with the same localisation pattern as the integrated strain, as shown by staining vacuoles of *NOP1*-GFP-Ktd1 cells and mitochondria of *CUP1*-GFP-Ktd1 and comparing expression levels (**Figure 6A-6C**).

**Figure 6:**
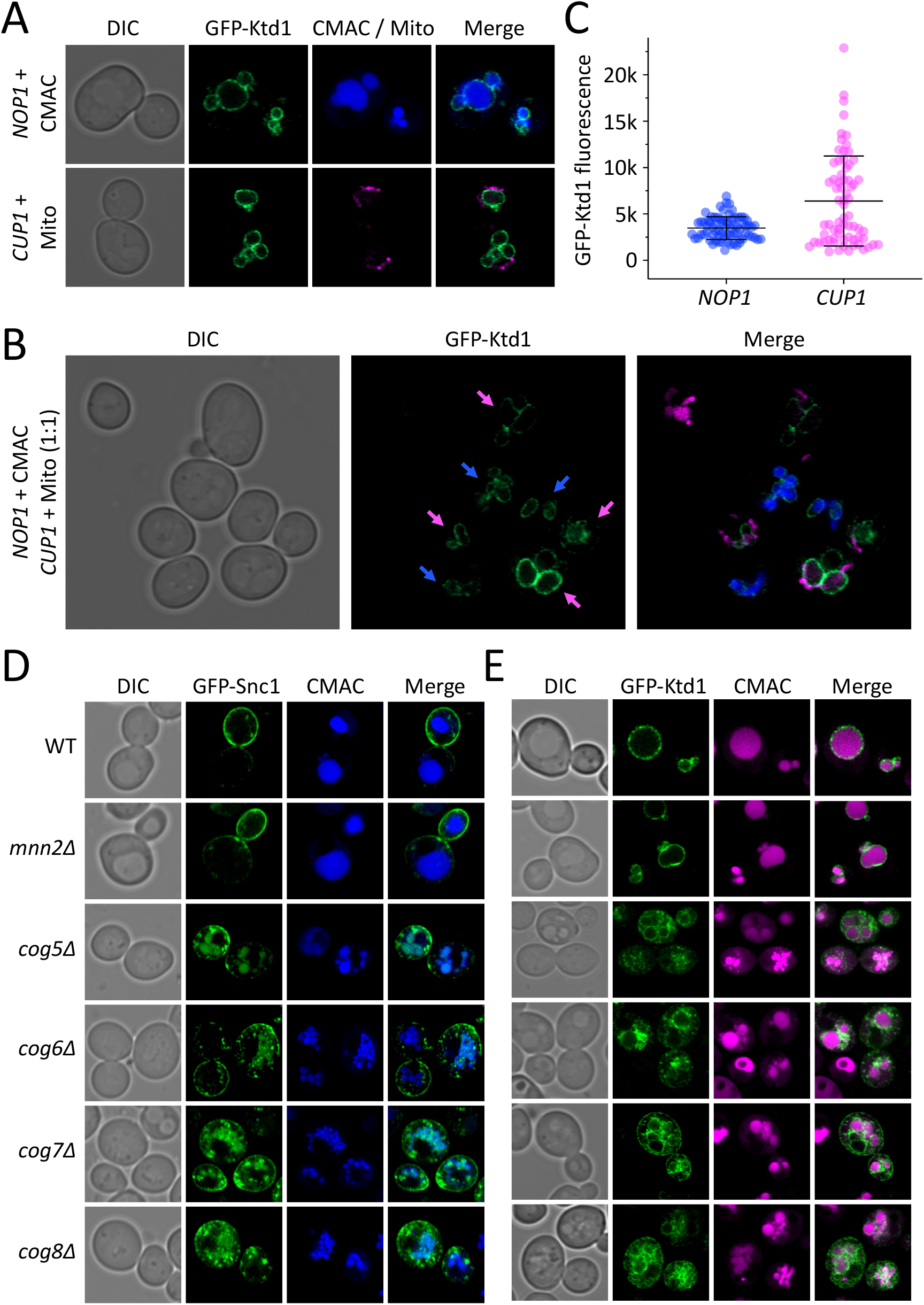
Correct localisation of Ktd1 requires the COG complex. **A - C)** *NOP1*-GFP-Ktd1 cells were stained with CMAC and wild-type cells expressing *CUP1*-GFP-Ktd1 from a plasmid were stained with MitoTracker Red. Cells were grown to mid-log-phase prior to labelling and washing, then prepared for confocal microscopy (**A**). The two cultures were mixed 1:1 and imaged together, blue (*NOP1*) and magenta (*CUP1*) arrows are indicated (**B**). GFP fluorescence from each condition was quantified by measuring integrated density of GFP signal from individual cells (**C**). **D)** Indicated strains expressing GFP-Snc1 (upper) and GFP-Ktd1 (lower) from a plasmid were grown to mid-log phase overnight, vacuoles were stained with CMAC (blue / magenta) prior to Airyscan2 confocal imaging. Error bars show standard deviation.

It has been previously shown that yeast synaptobrevin, a SNARE protein that localises mainly to the surface, is mis-localised in *cog* mutant cells (Whyte and Munro, 2001), likely as Snc1 recycles via the Golgi in a ubiquitin dependent manner (Laidlaw et al., 2022; Xu et al., 2017). We expressed GFP-Snc1 in *cog5Δ, cog6Δ, cog7Δ, cog8Δ* mutant strains and confirmed that all exhibit severe mis-localisation phenotypes, unlike wild-type and *mnn2Δ* cells, which correctly localise Snc1 to the surface in a polarised manner (**Figure 6D**). We note that in addition to various bright punctate GFP-Snc1 spots in the lobe B *cog* mutants there is also GFP signal in the vacuolar lumen. Indeed, there is also fragmented vacuolar phenotype indicated with CMAC staining in the mutant strains, which is reminiscent of vacuoles from *cog3 / grd20* mutants (Spelbrink and Nothwehr, 1999). As *cog* mutants disrupt trafficking of post-Golgi cargoes, we next asked whether they also mis-localise the membrane-spanning K28 toxin defence factor, Ktd1. As expected, GFP-Ktd1 localises to the vacuole membrane in wild-type and *mnn2Δ* cells. However, in lobe B *cog* complex mutants, GFP-Ktd1 has a severe mis-localisation phenotype (**Figure 6E**). As expression levels of GFP-Ktd1 do not change in these strains we assume this localisation phenotype is due to loss of COG-complex activity, and not an artifact of over-expression (**Supplemental Figure S2A**). Although some GFP-Ktd1 signal is found at the vacuolar membrane, there are many aberrant localisations, which includes large intracellular aggregates and signal at the endoplasmic reticulum. As the COG complex and Ktd1 are both required for K28 defence (**Figures 4A - 4D, Supplemental Figure S2B**) we conclude that the COG complex has a role in K28 toxin defence through coordinating precise endolysosomal trafficking of the membrane protein Ktd1.

### Ktd1 relies on the COG-complex for function

To explore the functional interplay between Ktd1 and the COG-complex we created mutants lacking *KTD1* in addition to each of the lobe B subunits of the COG complex (*COG5, COG6, COG7, COG8*). These combined mutations did not have a significant difference to the defect observed in *ktd1Δ* single mutants when measured with the halo assay (**Figure 7A**). Similarly, liquid based growth assays of *ktd1Δ cogΔ* combinations in media containing either K28 or the heat cured control all showed very similar sensitivities to the deletion of *KTD1* (**Figure 7B**). To assess K28 sensitivity of *cog* mutants exhibiting elevated levels of Ktd1, we first modified the diploid strain infected with the M28 virus, which is a uracil auxotroph due to its *ura3-52* / *ura3Δ* genotype and therefore not compatible with media used to select for over-expression plasmids. This was achieved by repairing the *URA3* locus and confirming the infected strain still effectively secreted K28 toxin (**Figure 8A - 8B**). We then transformed cells with either a vector control or *CUP1*-GFP-Ktd1, and performed halo assays using the Ura^+^ secretor strain. Interestingly, over-expressing GFP-Ktd1 in a wild-type background provides additional resistance to K28, in addition to functionally complementing *ktd1Δ* cells (**Figure 8C**). Whilst some additional protection against K28 was also observed in lobe B *cogΔ* cells, the relative levels of rescue were much less than was seen in WT cells. This would be explained by most of the additional Ktd1 being non-functional in *cogΔ* mutants due to mislocalisation.

**Figure 7:**
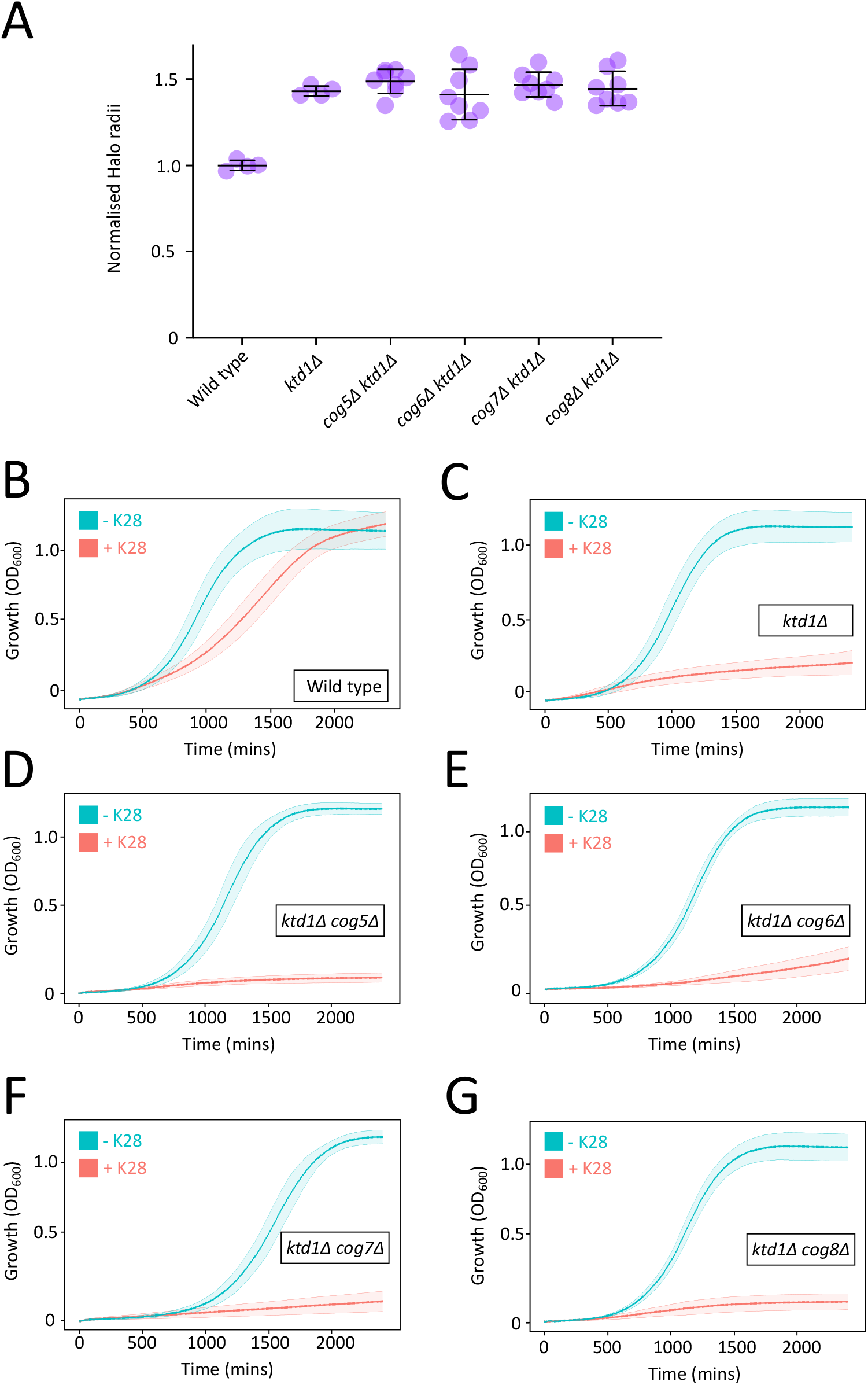
Combining *ktd1* and *cog* mutations does not increase K28 sensitivity. **A)** Halo assays normalised to a wild-type control were compared for *ktd1Δ* single and double *cogΔ* mutants. Error bars show standard deviation. **B - G)** The indicated yeast strains were assessed for growth in the presence (pink) and absence (blue) of K28, with graphs generated using R:Shiny app described in Figure 3.

**Figure 8:**
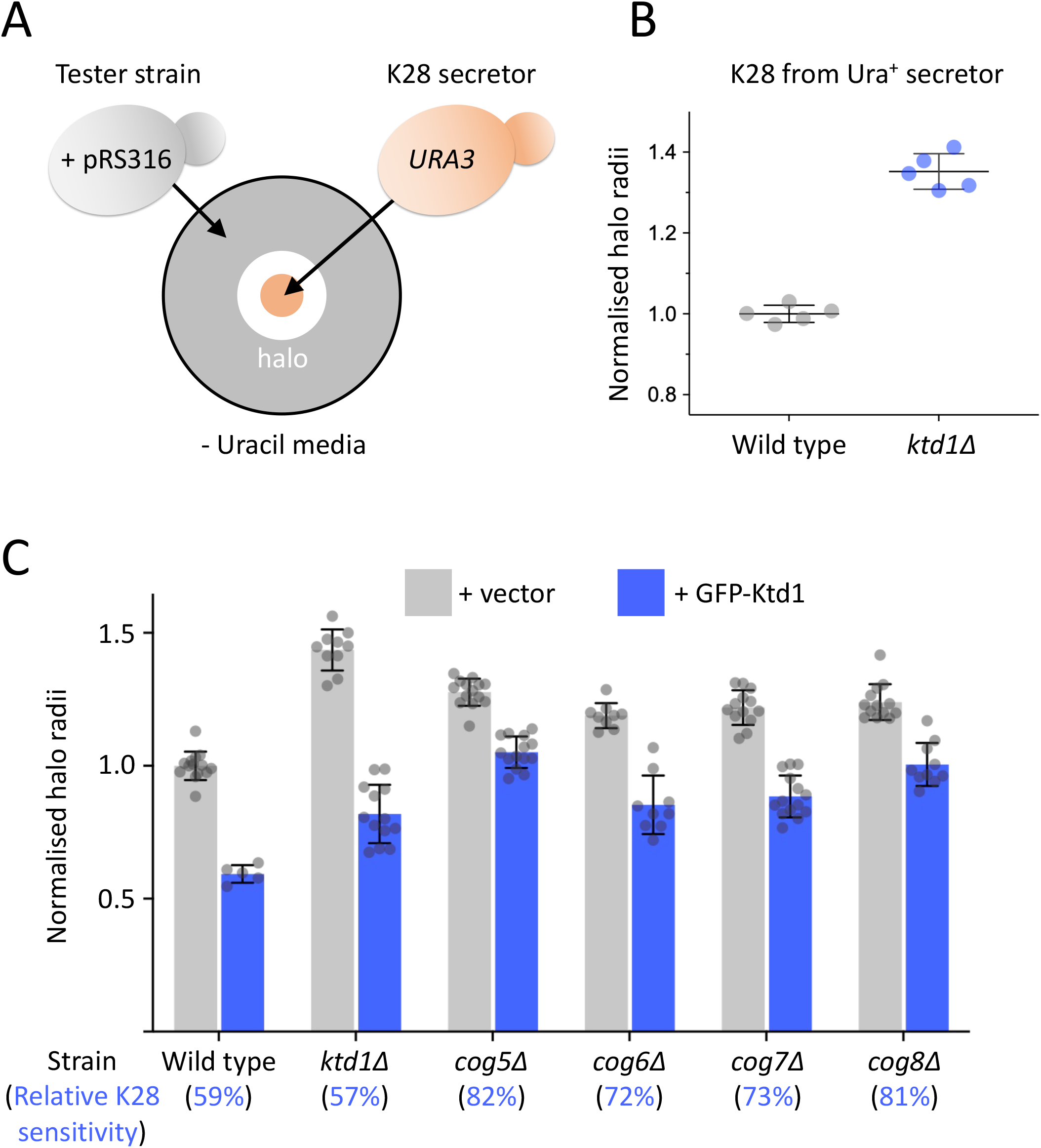
Over-expression of Ktd1 has reduced capacity to defend against K28 in *cog* mutants. **A)** Schematic diagram showing the requirement of K28 secretor strains to be prototrophic for uracil, by repairing the *URA3* gene, to allow halo assays to be performed in SC-uracil media that can select for tester lawn strains to harbour a pRS316-based *URA3* plasmid. **B)** The Ura^+^ converted K28 secretor strain was used in halo assays for wild-type and *ktd1Δ* mutants on SC media. Error bars show standard deviation. **C)** Indicated strains were transformed with empty pRS316 vector (grey) or the *URA3* based *CUP1*-GFP-Ktd1 plasmid (blue) before halo assays were performed in SC-uracil media. Halo radii normalised to wild-type + vector are shown, with the percentage of additional rescue to K28 defence, compared to vector control, shown in parenthesis.

## DISCUSSION

This study investigates the role of the COG complex in yeast resistance to the Killer Toxin 28 (K28) and provides evidence that its function is essential for proper trafficking of the K28 defence factor Ktd1 to post-Golgi endolysosomal compartments. Our findings extend the current understanding of how intracellular trafficking contribute to cellular resistance against toxins of the A/B family.

Our study confirms and expands upon previous observations implicating the COG complex in K28 toxin sensitivity. Specifically, we show that deletion of non-essential lobe B subunits of the COG complex (Cog5–8) results in a significant increase in K28 sensitivity. This suggests that the functional integrity of the entire lobe B subcomplex is critical for toxin defence and may have unique functional role in K28 defence that is distinct from the canonical functions of the COG complex. Alternatively, the entire COG complex is required for mediating efficient defence against K28 and the essentiality of the lobe A mutants make functional testing of those subunits uninformative; we speculate the latter is most likely. We tested various mutant alleles of the essential lobe A subunits, with reduced levels of different subunits, but these failed to show phenotypes when tested at elevated temperature or following exposure to K28 (**Supplemental Figure S1**). While prior studies identified Cog7 as being important for K28 resistance (Carroll et al., 2009), our data demonstrate that the other lobe B subunits are equally critical. We speculate that although the previous genetic screen was performed in an extremely robust manner, with mutants being validated extensively, the screen suffered from the relatively low resolution ‘halo’ based assay, which is particularly difficult to perform at scale.

This prompts considerations of assays for K28 sensitivity. Enriching media with K28 from a diploid hyper-secreting strain, with control media from heat-cured cells that do not secrete K28, is effective in liquid assays (Andreev et al., 2023; Schmitt and Tipper, 1990). Performing these liquid growth measurements at 96-well density increases capacity and our bespoke analysis package generates extensive data, including growth profiles ± K28 toxin and comparisons to reference selected strains (e.g. wild-type cells). However, we also note that wild-type cells - which endocytose and tolerate K28 via Ktd1 - do not have capacity for significantly better growth to entirely resistant strains. Although the halo-based K28 assay does have a small dynamic range to distinguish sensitive, resistant or wild-type phenotypes, we found it very effective at highlighting the differences between wild-type cells and resistant strains, such as those lacking mannosyltransferases (**Figure 3D**). We recommend using both assays in the first instance. Our optimisation of the halo assay also allows for conditions that at least provide the best range for comparisons. We assume slower growth on the lawn in SC media allows toxin to permeate more efficiently, whilst the secretor strain either grows or secrete more effectively when grown in rich media.

A detailed structure of the target glycan epitope for K28 is unknown, however, based on the strong resistance phenotypes of the *mnn1Δ, mnn2Δ*, and *mnn5Δ* mutants (Carroll et al., 2009), it is likely that K28 binds to terminal 1,2 and/or 1,3 linked mannose residues. Defects in the COG complex, which results in mis-localisation or degradation of Golgi localised glycosyltransferases, have an impact on glycosylation profiles at the yeast cell surface (Smith and Lupashin, 2008). However, adsorption assays revealed no significant difference in K28 binding between wild-type and all lobe B *cog* mutant cells, and double mutants lacking both COG subunits and *MNN9* exhibited resistance similar to *mnn9Δ* single mutants (**Figure 5B**). These results suggest that the hypersensitivity of *cog* mutants to K28 is not caused by creating an alternate route to adsorption. We next hypothesised that the increased K28 sensitivity in *cog* mutants could result from defects in intracellular trafficking rather than glycosylation. GFP-Snc1 is known to recycle via the Golgi with steady state localisation at the surface reliant on efficient recycling via endosomally localised COPI and other factors, including the COG complex for proper sorting to post-Golgi compartments (MacDonald and Piper, 2016; Whyte and Munro, 2001). Similarly, we observed severe trafficking defects of GFP-Ktd1 in *cog5Δ, cog6Δ, cog7Δ, and cog8Δ* mutants. Unlike wild-type cells, where Ktd1 localises predominantly to the vacuolar membrane, *cog* mutants exhibit aberrant Ktd1 localisation, including bright intracellular puncta and signal inside the vacuolar lumen (**Figure 6**). The idea that defects in K28 defence in *cog* mutants is directly explained by Ktd1 is supported by the combination of these mutations in haploid strains having no additive effects (**Figure 7**), although we acknowledge the defects of *ktd1Δ* are very severe and do not leave a large dynamic range for additional sensitivity to be observed. Curiously, Snc1 sorting to the vacuole requires the Cos proteins from the *DUP380* family (MacDonald et al., 2015a; MacDonald et al., 2015b), which are evolutionarily related to Ktd1 from the *DUP240* family (Despons et al., 2006).

The steady state mis-localisation phenotype of GFP-Snc1 in *cog* mutants is compounded by multiple rounds of surface recycling via the Golgi (Lewis et al., 2000). One attractive idea to explain why Ktd1 is so affected by the *cog* mutations is that it too cycles via the *trans*-Golgi network multiple times, and not simply a single Golgi pass *en route* to the vacuole. This hypothetical model is based on the fact that Ktd1 localises to early endosomes, multivesicular bodies and the *trans*-Golgi network (Andreev et al., 2023). If Ktd1 routinely transits these compartments as a toxin surveillance mechanism upstream of its terminal destination at the vacuolar membrane, one would predict a severe mis-localisation upon disruption of the required membrane trafficking machinery. This is further supported by the observation that over-expression of GFP-Ktd1 in *cog* mutants cannot effectively rescue toxin defence to the levels of wild-type or *ktd1Δ* mutants, suggesting extra levels of Ktd1 are also mislocalised and non-functional (**Figures 8**). We do not exclude the possibility that the COG complex also has a role in trafficking K28 in a protective mechanism that does not relate to Ktd1.

The mis-localisation of Ktd1 in *cog* mutants provides a mechanistic explanation for their hypersensitivity to K28. Ktd1 is a key defence factor that localises to the vacuolar membrane, where it likely functions to neutralise internalised K28 toxin or prevent its retrograde trafficking. Disruption of Ktd1 localisation in *cog* mutants would impair its protective function, rendering cells more susceptible to K28-induced cell death. The requirement for the COG complex in Ktd1 trafficking underscores the importance of membrane trafficking pathways in cellular defence against such toxins. We speculate that COG facilitates Ktd1 sorting between different endosomal sub-compartments, most likely by mediating its cycling through the TGN. This sorting ultimately culminates in accumulation of Ktd1 at the vacuole. Therefore, when K28 internalises to endosomes surveyed by Ktd1, Ktd1 might be able to engage toxin and divert it to the vacuole instead of retrograde trafficking. The findings presented here have broader implications for understanding cellular resistance mechanisms against A/B toxins. Lysosomes in animal cells swell in response to endocytosed toxin (Turner et al., 2023) and lysosomal degradation of Shiga toxin occurs if retrograde trafficking to the Golgi is blocked (Mukhopadhyay and Linstedt, 2012); therefore, it is possible that analogous mechanisms involving the trafficking of defence factors may exist in other eukaryotes.

## MATERIALS & METHODS

Information about yeast strains and plasmids used in this study can be found in supplemental table 1. Relevant statistical test results are included in in supplemental table 2.

### Cell culture

Yeast cultures were typically grown in rich yeast extract peptone dextrose (YPD) medium 2% glucose, 2% peptone, 1% yeast extract or synthetic complete (SC) minimal medium 2% glucose, yeast nitrogen base supplemented with appropriate amino acids and bases. For making agar plates, 2% agar is also added prior to autoclaving. Cells were routinely grown overnight at 30°C with shaking to early/mid-log phase (OD_600_ ≤ 1.0) prior to experimentation.

### Plate based growth assays

Yeast cultures were grown to log phase in SC or YPD media as indicated, before OD_600_ measurements were taken and used to normalise cultures, and then used to create 10-fold serial dilutions in a microtiter plate. Yeast cultures were then spotted on agar media plates, allowed to dry at room temperature and then incubated at indicated temperatures. Growth of plates was then recorded, typically after 24 - 48 hours.

### Halo K28 sensitivity assay

Strains for assessment were cultured to saturation overnight, then OD_600_ measurements were used to normalise lawn densities to OD_600_ = 0.1 - 0.2, unless otherwise stated. Lawns were plated on agar media containing phosphate citrate buffer adjusted to pH 4.7. A separate culture of the *ski2Δ*/*ski2-2* diploid strain infected with the M28 virus was cultured to log phase (OD_600_ ∼ 1.0) before concentration 10-fold and 5 - 7 technical repeats per plate spotted on to the lawns. Yeast were grown for 48 hours at room temperature before growth was recorded. Images were contrast adjusted using Fiji / ImageJ and the radii of halos were measured with 4 individual measurements per spot across all technical replicates and data from biological replicates plotted.

### Purification of enriched media

K28 toxin secretor cells and a heat-cured negative control strain were grown in SC media pH 4.7 for 48 hours at 23°C with mild agitation at 70 rpm. The K28 enriched media was then filter sterilised and stored on ice, before addition of 5x concentrated SC media (pH 4.7) to replenish nutrients. Growth of cells was measured using the Clariostar (BMG LabTech) automated plate reader set to absorbance at 600 nm in a 96-well plate format. Wells were prepared with 4 - 6 technical replicates, containing 200μl of K28 toxin containing, or the heat cured control media. Strains for assessment were cultured to saturation overnight, then ∼1.5 μl added to each well. Absorbance measurements were taken every 10 minutes for 27 - 32 hours. Prior to each measurement, the plate underwent orbital shaking for 30 seconds. Raw data was then organised and read with an R:Shiny Package (made available: https://shiny.york.ac.uk/bioltf/cm_growthcurves/) to generate graphs and statistics. The coding and scripting information related to this application have been housed in the GitHUb repository, details can be foun d here: https://github.com/uoy-research/cm_growthcurves.

### K28 adsorption assays

To test K28 binding to different strains, toxin enriched media was processed as described above before. Strains to be tested were cultured overnight and OD_600_ used to harvest equal amounts of each strain. Pellets were incubated for 15 minutes at 4°C in toxin enriched media, with all comparisons in each biological replicate performed at the same with the same media preparation. Following the adsorption period, cells were removed and the resultant media filtered again and then used in growth assays, also described above, to assay the efficacy of any remaining toxin in the media. Area under the curve following 32 hours growth was normalised to wild-type controls on the day, and then plotted as an indirect measure of toxin in the media.

### Confocal microscopy

Wild type and knock out yeast strains expressing GFP-tagged proteins were grown overnight to mid-log phase, and imaged at room temperature using a 63x 1.40 oil immersion Plan Apochromat objective lens on a Zeiss LSM 980 with Airyscan2. GFP fluorescence was excited using a 488 nm Argon laser and 495–550 nm emission collected. Yeast vacuoles were labelled with 50 μM 7-amino-4-chloromethylcoumarin (CMAC) and washed x3 prior to confocal imaging. Blue fluorescence of CMAC was excited with a 405 nm laser and emissions collected at 422/497 nm. 2 μM MitoTracker Red CMXRos (Thermo) was added to cultures for 30 minutes, prior to washing and imaging, with excitation using a 594 nm laser and emissions 570/620 nm. Processed images were adjusted in Fiji/ImageJ software (NIH).

## Supporting information

Combined PDF with table legend sand 3 supp figures

Reagent table

Stats

## ACKNOWLEDGMENTS

We would like to thank staff at the York Bioscience Technology Facility for technical assistance. This research was supported by a Sir Henry Dale Research Fellowship from the Wellcome Trust and the Royal Society 204636/Z/16/Z (CM). This research was also supported in part by the Intramural Research Program of the National Human Genome Research Institute, National Institutes of Health (IA and MJS).

## CONTRIBUTIONS

Conceptualization: IA, DU, MS, CMD

Methodology: MS, CMD

Software: AD, SM

Validation: KL, HN, AM, MX

Investigation: KL, HN, AM, MX, JS, CMD

Writing - Original Draft: CMD

Writing - Review & Editing: HN, AM, CMD

Resources: AL

Visualization: KL, AD, SM, CMD

Supervision: DU, CMD

Project administration: CMD

Funding acquisition: CMD

## FIGUREs & LEGENDS LEGENDS

**Supplementary Figure S1:**
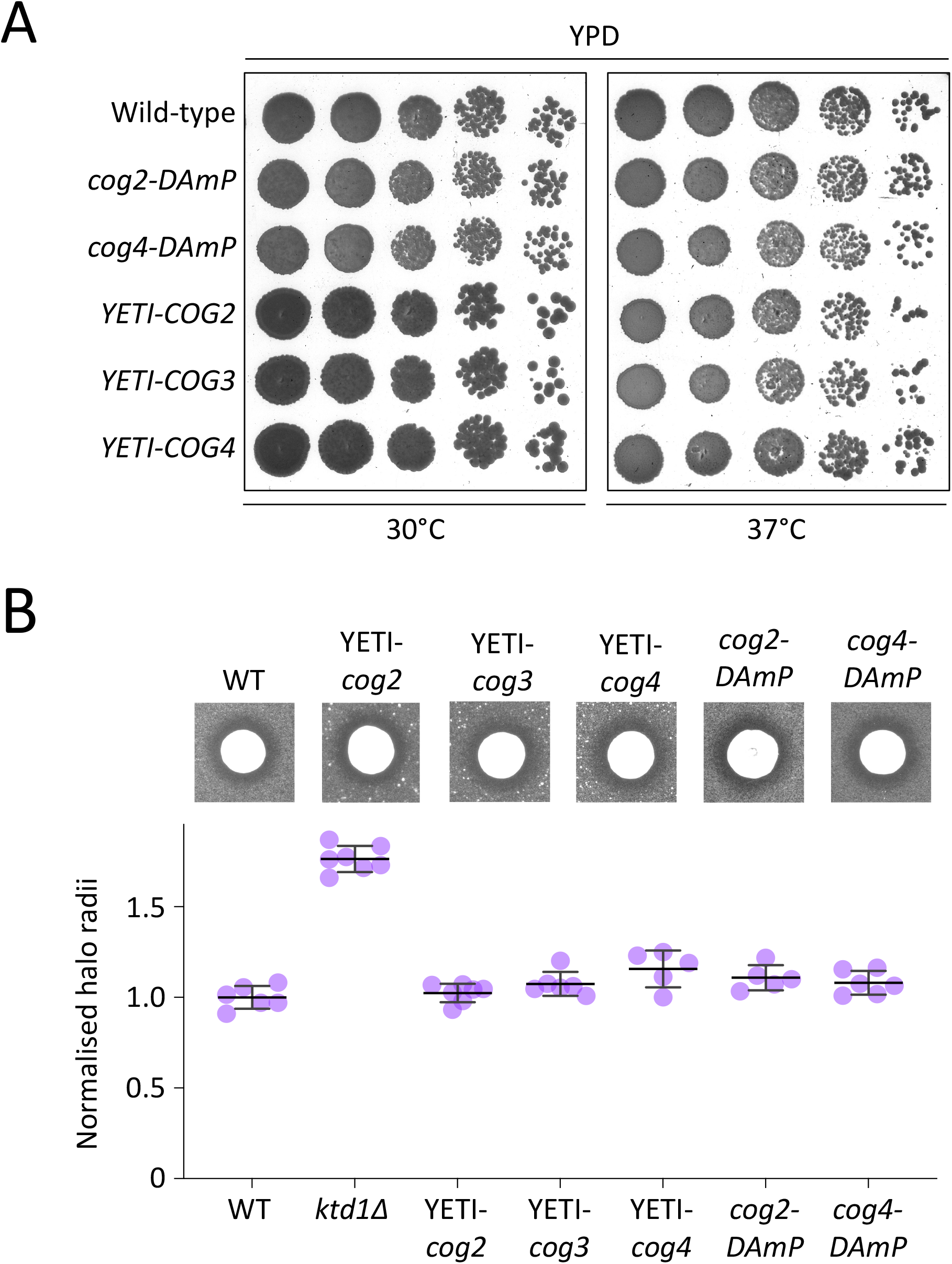
Testing lobe A mutants of the COG complex. **A)** Wild-type (WT) and indicated mutants were cultured in YPD media to mid-log phase before normalised cell amounts were harvested and used to create a 10-fold serial dilution and plating on to YPD agar before incubation at either 30°C or 37°C prior to recording growth. **B)** K28 sensitivity halo assays were performed using indicated strains (above) with quantified halo radii (n > 4) plotted (below). Error bars show standard deviation.

**Supplementary Figure S2:**
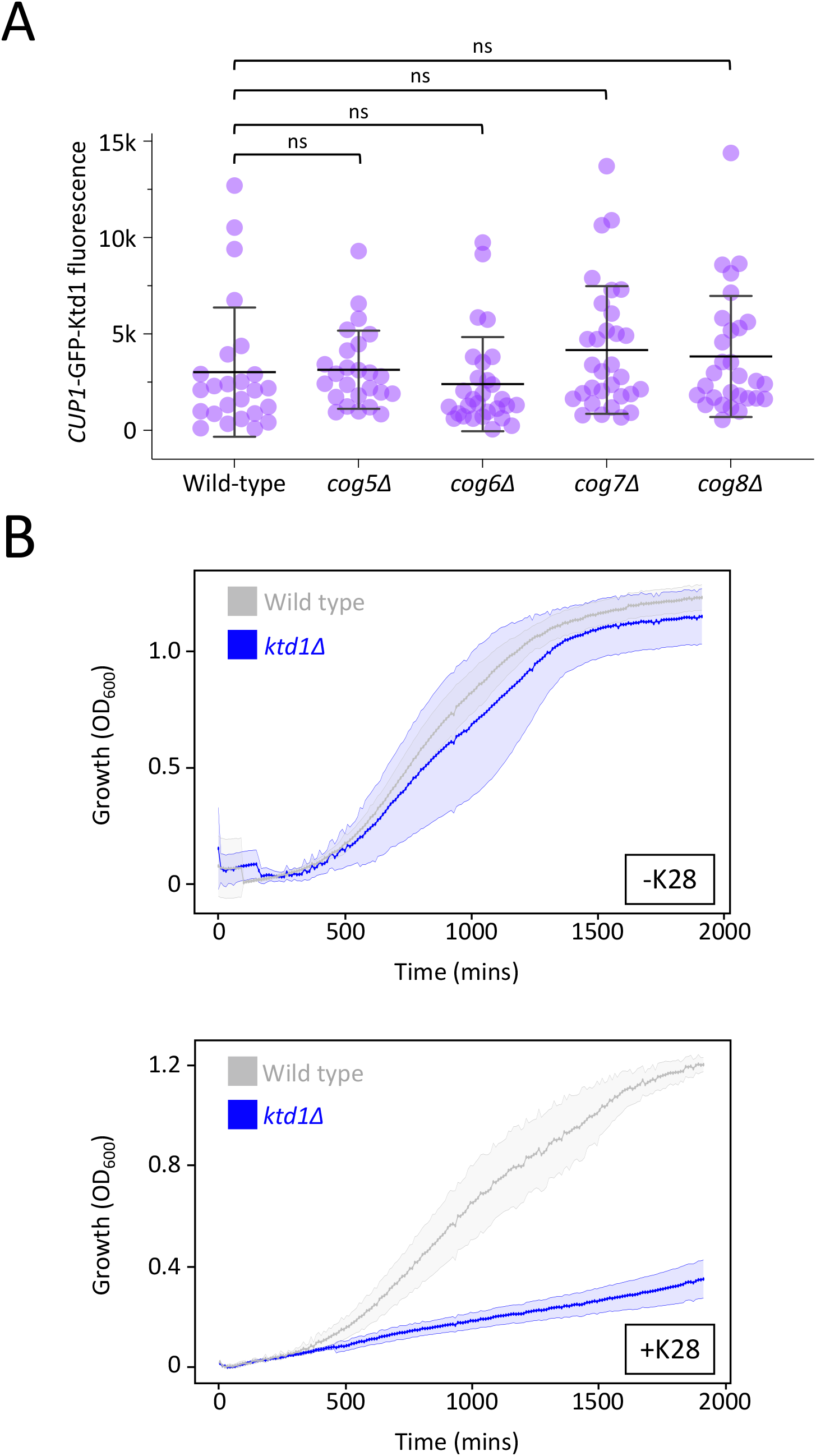
Expression and functional analyses of Ktd1. **A)** A plasmid expressing GFP-Ktd1 from the *CUP1* promoter in media lacking additional cooper was compared across indicated strains, with integrated density of GFP fluorescence measured per cell and plotted. **B)** Growth curves comparing wild-type (grey) and *ktd1Δ* mutants in control media (upper) and active K28 containing media (lower). Error bars show standard deviation.

**Supplementary Figure S3:**
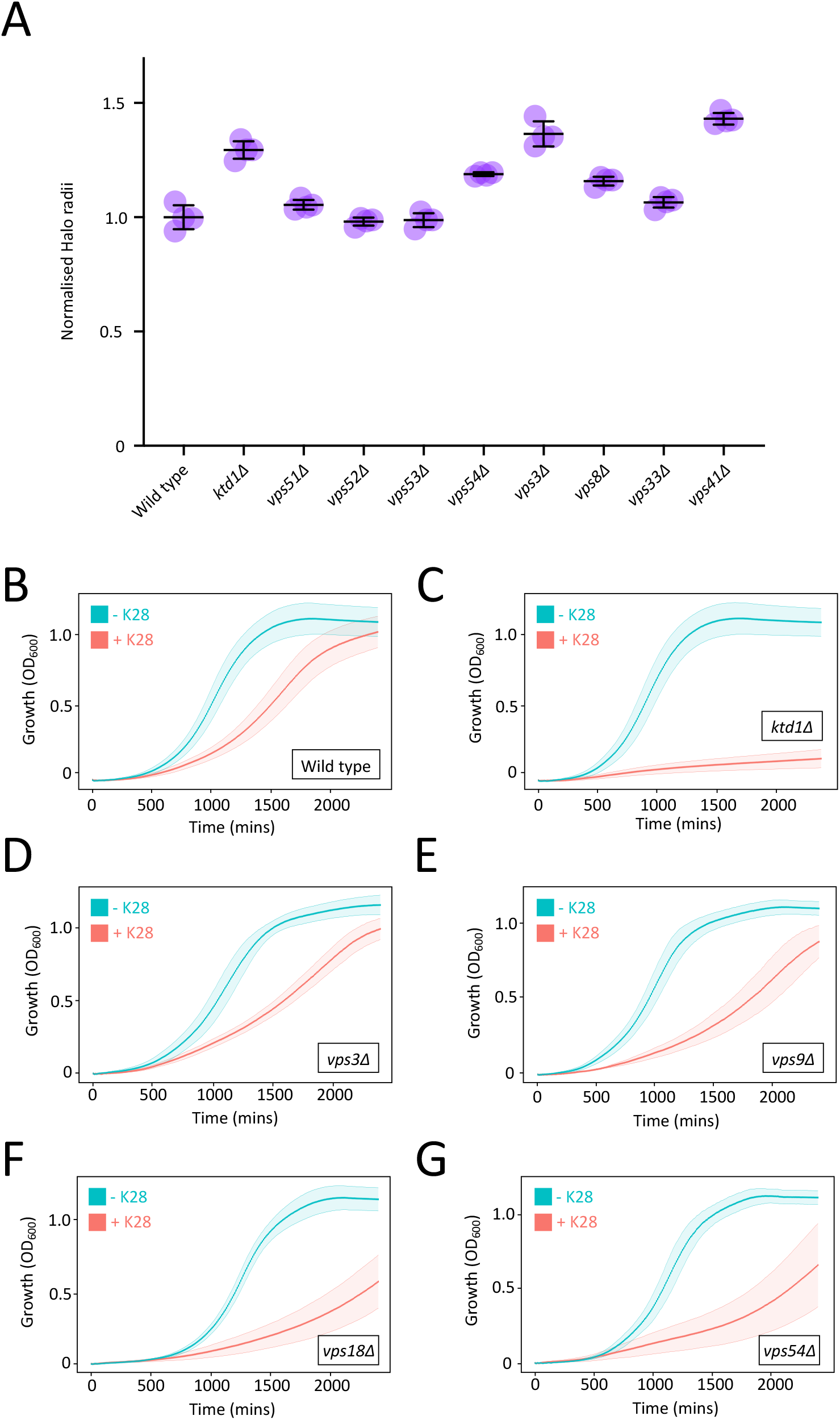
K28 sensitivity measurements in additional trafficking complex mutants. **A)** K28 sensitivity halo assays of indicated mutants were performed in SC media and normalised to wild-type controls. **B - G)** Liquid groth assays for indicated yeast strains grown in K28 (pink) or heat-cured control (blue) media.

