## Supplementary material for "Killer Toxin K28 resistance in yeast relies on COG complex mediated trafficking of the defence factor Ktd1": Combined PDF with table legend sand 3 supp figures

### SUPPLEMENTAL TABLE LEGENDS

### SUPPLEMENTAL FIGURES & LEGENDS

**Supplemental Table S1: Reagent table**
Table listing all the strains and plasmids used in this study, including database and source information and other relevant information.

**Supplemental Table S2: statistical tests**
Table listing statistical tests and comparisons performed, including reference to the sub-panel / experiment from the manuscript.

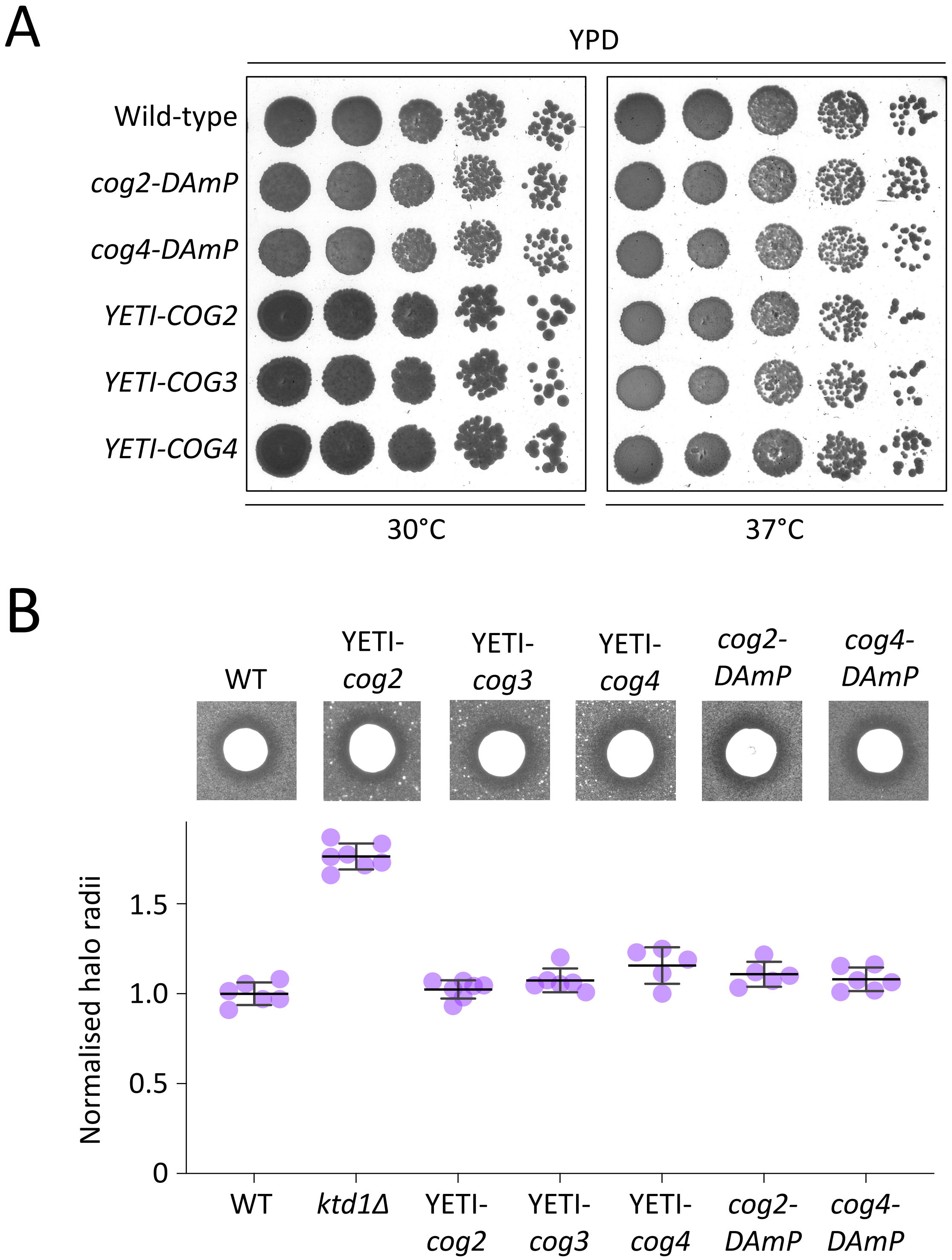


**Supplemental Figure S1: Testing lobe A mutants of the COG complex**
**A)** Wild-type (WT) and indicated mutants were cultured in YPD media to mid-log phase before normalised cell amounts were harvested and used to create a 10-fold serial dilution and plating on to YPD agar before incubation at either 30°C or 37°C prior to recording growth.
**B)** K28 sensitivity halo assays were performed using indicated strains (above) with quantified halo radii (n > 4) plotted (below). Error bars show standard deviation.

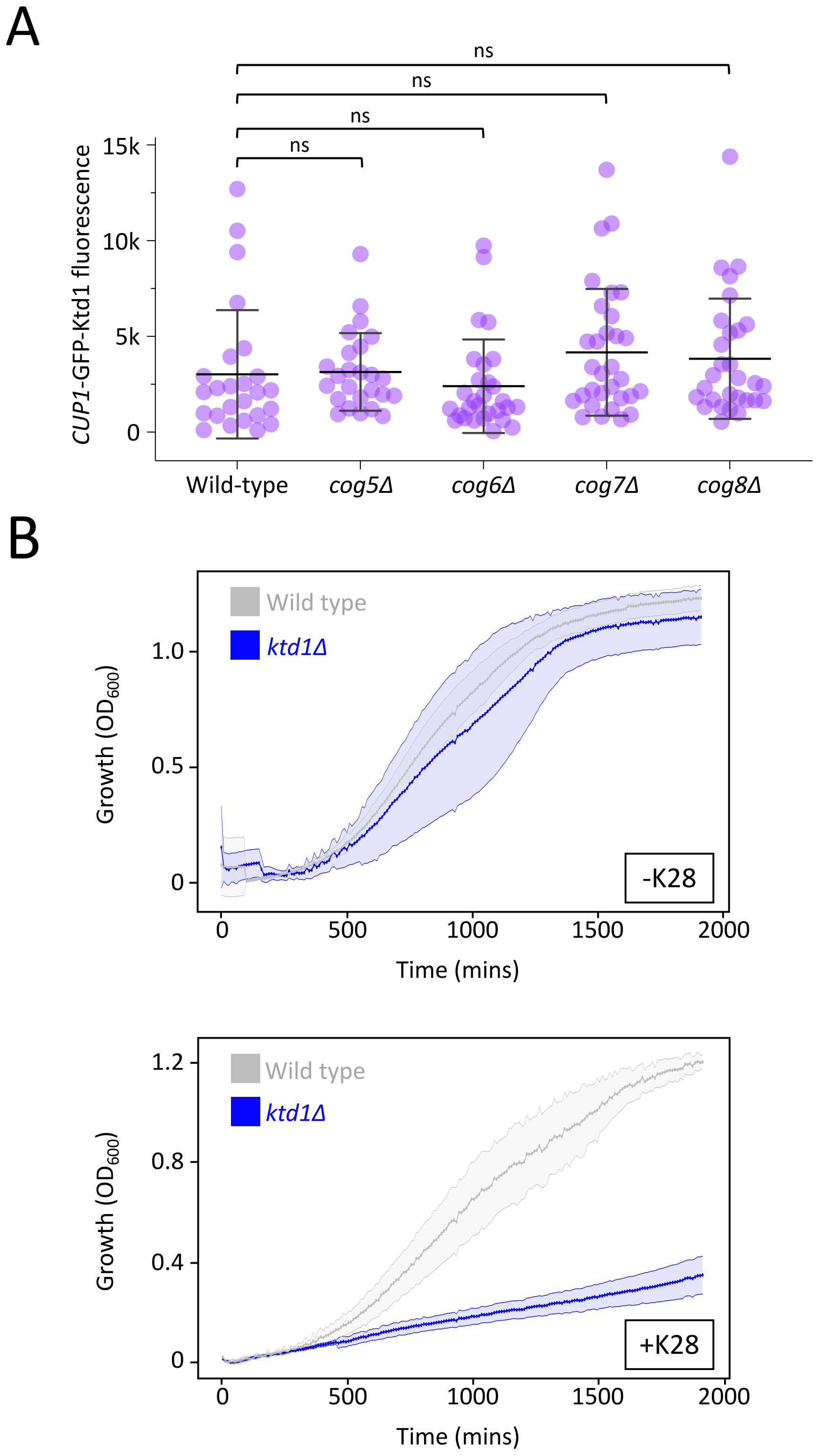


**Supplemental Figure S2: Expression and functional analyses of Ktd1**
**A)** A plasmid expressing GFP-Ktd1 from the *CUP1* promoter in media lacking additional cooper was compared across indicated strains, with integrated density of GFP fluorescence measured per cell and plotted. **B)** Growth curves comparing wild-type (grey) and *ktd1*Δ mutants in control media (upper) and active K28 containing media (lower). Error bars show standard deviation.

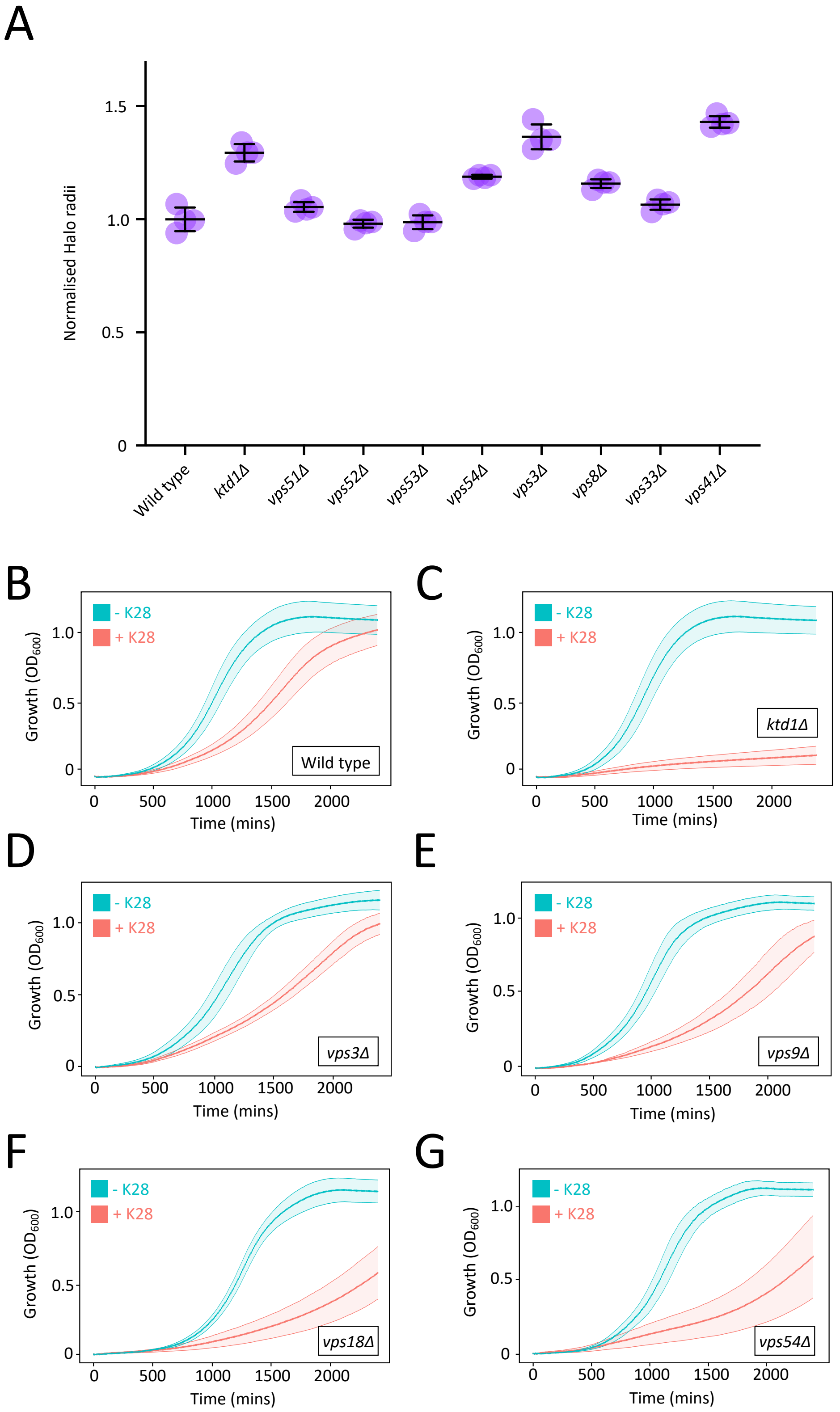


**Supplemental Figure S3: K28 sensitivity measurements in additional trafficking complex mutants**
**A)** K28 sensitivity halo assays of indicated mutants were performed in SC media and normalised to wild-type controls.
**B - G)** Liquid growth assays for indicated yeast strains grown in K28 (pink) or heat-cured control (blue) media.
